# Deciphering programs of transcriptional regulation by combined deconvolution of multiple omics layers

**DOI:** 10.1101/199547

**Authors:** Daniel Hüebschmann, Nils Kurzawa, Sebastian Steinhauser, Philipp Rentzsch, Stephen Krämer, Carolin Andresen, Jeongbin Park, Roland Eils, Matthias Schlesner, Carl Herrmann

## Abstract

Metazoans are crucially dependent on multiple layers of gene regulatory mechanisms which allow them to control gene expression across developmental stages, tissues and cell types. Multiple recent research consortia have aimed to generate comprehensive datasets to profile the activity of these cell type- and condition-specific regulatory landscapes across many different cell lines and primary cells. However, extraction of genes or regulatory elements specific to certain entities from these datasets remains challenging. We here propose a novel method based on non-negative matrix factorization for disentangling and associating huge multi-assay datasets including chromatin accessibility and gene expression data. Taking advantage of implementations of NMF algorithms in the GPU CUDA environment full datasets composed of tens of thousands of genes as well as hundreds of samples can be processed without the need for prior feature selection to reduce the input size. Applying this framework to multiple layers of genomic data derived from human blood cells we unravel mechanisms of regulation of cell type-specific expression in T-cells and monocytes.

## Background

Multicellular organisms heavily rely on the specialized function of various tissues, each of which is composed of a multitude of distinct cell types. To achieve this great intra-organismal complexity from cells that all carry the same DNA sequence, multiple layers of gene expression regulation cooperate to tune transcript and ultimately protein levels and promote distinct cellular phenotypes [1]. Regulatory elements which are clusters of transcription factor binding sites can regulate expression of nearby and remote genes [2]. Thereby, especially distal acting regulatory elements such as enhancers are crucial to establish and maintain cell type-specific gene expression [3].

In recent years, several large consortia attempted to understand cellular gene expression regulation by generating huge datasets comprising multiple assays and cell types [4–7]. However, it is fay beyond trivial to discover features (e.g. genes or genomic loci such as enhancers) in these datasets that specifically correspond to one unique entity which can be different tissues, developmental stages or different treatment conditions or features that contribute to distinct processes or developmental stages [8].

Principal component analysis (PCA) is commonly used to explore high-dimensional datasets and can be exploited to extract features that separate certain subgroups. However, due to PCA‘s freedom of mixing negative and positive feature contributions, it does not necessarily resolve fine differences between entities in big datasets in a sufficient manner. In contrast to PCA, non-negative matrix factorization (NMF) which aims at finding two non-negative matrices that factorize an input matrix, forces all features to be represented in an additive manner. Hence NMF leads to a “parts-based representation”, and thus to a better interpretability of the derived signatures [9]. NMF has been used in different settings for analysis of Next Generation Sequencing (NGS) data [10–21]. In this article, we introduce a novel framework implemented in R called Bratwurst, which allows efficient matrix decomposition of any genomic data type represented as a numeric matrix. We take advantage of implementations of NMF algorithms in the GPU CUDA environment. Given the performance of these GPU implementations, full datasets composed of tens of thousands of genes or hundreds of thousands of genomic regions, as well as hundreds of samples can be processed, without the need for prior feature selection to reduce the input size.

Besides the NMF algorithm itself, we present a novel strategy to combine omics datasets by comparing and matching signatures originating from distinct data types. Using different pre-labeled cell types of the human hematopoietic system [22], we show how this method can be used to achieve a dimensionality reduction, to produce a powerful visualization of the data and to extract cell type-specific features.

## Results and Discussion

### A novel workflow for GPU-based decomposition of genomic datasets

We implemented the R package Bratwurst, which provides a multi step workflow for applying GPU-based NMF to genome-wide signals across multiple omics layers (available at https://github.com/wurst-theke/bratwurst). In addition, Bratwurst includes functionalities to determine the optimal factorization ranks and to extract signature-specific features. The workflow is schematically illustrated in Figure 1:

**Figure 1:**
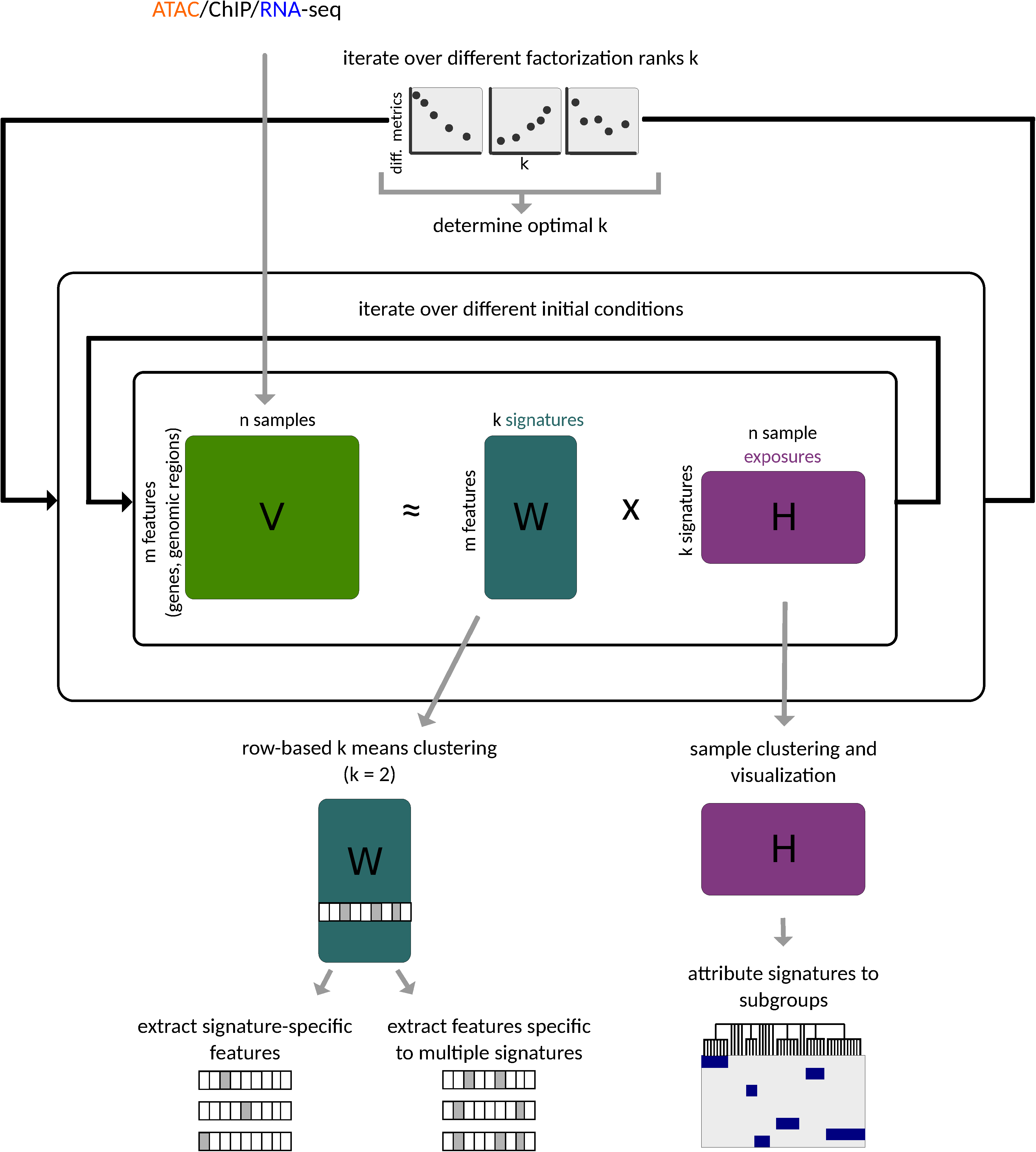
Schematic illustration of our framework for performing non-negative matrix factorization and downstream analysis. The factorization is performed for a number of different consecutive *k* and different initial conditions. After an optimal factorization rank is determined the matrix *H* is used for clustering and visualization of samples and thus interpretation of the signatures. The retrieved matrix *W* is used for feature extraction of the different signatures.

- **Step 1: Preprocessing**. Before running an NMF analysis, data pre-processing is highly recommended. Emphasis should be placed on two aspects: (i) reduction of sparsity of the input matrix (e.g. by removing rows and columns which have only zero entries or entries smaller than a user-defined cutoff) and (ii) centering and rescaling of the data (e.g. with the help of the function `estimateSizeFactorsForMatrix()` from the R package DESeq2 [23]). As numerous techniques for scaling, centering and filtering of data are available, the workflow offers a wide flexibility to the user at this step.
- **Step 2: NMF**. NMF is a family of algorithms to factorize one large matrix *V* (of dimensions *n* x *m*) into two smaller matrices *W* (the *signature matrix* of dimensions *n* x *k*) and *H* (the *exposure matrix* of dimensions *k* x *m*) under the constraint of non-negativity on all entries in both factor matrices *W* and *H* (Figure 1). *k* is called the *factorization rank*; a complexity reduction is achieved if *k* < *n* and *k* < *m*. In addition to a novel CUDA-based NMF implementation, the Bratwurst package allows to use different existing NMF implementations like the CUDA-based NMF_GPU [24] or a CPU implementation from the R package NMF [25]. The factorization rank *k* is a free parameter for any NMF method. The optimal factorization rank can be determined with the following strategy [26]: iterate over different factorization ranks, evaluate quality metrics for any one of them and choose the rank which complies best with the quality metrics. Relationships between signatures extracted at different factorization ranks *k* may be visualized as Sankey diagrams or riverplots [27] (Figures S3 and S4). For details we refer to the methods section.
- **Step 3: Mapping signatures to labelled groups of samples**. The exposures (encoded in the matrix *H*) represent the contributions of the signatures to the different samples. These are used to group the samples and attribute subgroups to signatures. Names for the signatures are assigned generically by the function `getSignatureNames()` based on available group labels provided by the user. The assignment of subgroup names and other properties (e.g. colour coding) to the extracted signatures can be transferred to the riverplot visualization (Figure S3 and S4), showing at which factorization rank which pattern arises.
- **Step 4: Feature selection**. Once signatures are assigned to subgroups of samples, features with a high contribution to a specific signature can be extracted using the following non-parametric procedure:
  - (i) For every feature *f*, i.e. every row of the matrix *W*, perform a k-means clustering with *k* = 2. In general, there will be a subset of signatures with high contributions of *f* and a subset of signatures with low contributions of *f*.
  - (ii) For each feature, we obtain a binary vector across the signatures indicating to which of the signatures this feature has particularly high contributions. We can now find features which contribute highly to one single signature (**single-signature features)**. Conversely, features which contribute substantially to more than one signature are called **multi-signature features**.
  The extracted single-signature or multi-signature features can be used for downstream analyses e.g. enrichment analyses (see Methods).

For datasets with multiple data layers, multi-omics integration can now be performed by analyzing every layer individually (steps 1-4) and relating the resulting signatures by pairwise comparisons.

In the next section, we will present an example of such a multi-omics integration based on matching expression and chromatin accessibility datasets obtained in human blood cells.

### Transcriptional and regulatory programs in hematopoiesis

We applied our integrative approach to a recently published dataset of sorted blood cell populations, for which both RNA-seq and ATAC-seq are available [22]. In this work, we used a subset of 12 cell populations across 45 samples for RNA-seq and 68 samples for ATAC-seq, covering different branches of the haematopoietic tree, and various stages of differentiation from stem cells to terminally differentiated cells. For 24 samples, both data types were available.

Applying the NMF workflow on both datasets, the optimal factorization ranks k_RNA-seq_= 8 and k_ATAC-seq_= 9 were found. Supplemental Figure S1 illustrates different metrics justifying the choice of these factorization ranks. The choice of the optimal factorization rank is highlighted by transparent grey rectangles. Remarkably these distinct datasets yielded very similar optimal factorization ranks, i.e. very similar numbers of signatures, suggesting that there is a common underlying structure relating the regulatory landscape of open chromatin regions to the transcriptional landscape. For both decompositions, most signatures were highly specific to a particular cell population or group of cell populations (Figures 2a and 2c). Signatures extracted from RNA-seq were labeled R1 - R9 (note that there is no R2, therefore this corresponds to eight labels) and signatures extracted from ATAC-seq were labeled A1 - A9. For example, signature R5 of the RNA-seq data showed a high specificity for CD4+/CD8+ T-cells, while signatures R7 and R8 were specific to the NK cells and monocytes, respectively. Gene ontology enrichment analyses with the R package topGO [28]of the single-signature features extracted from the expression signatures revealed GO terms matching the biological function of the respective cell types: for the T-cell expression signature many terms related to T-cell differentiation, activation and co-stimulation, for the monocyte expression signature terms including complement receptor activity and opsonin receptor activity and for the NK cell expression signature terms like natural killer cell mediated immunity, cytotoxicity, chemotaxis, innate immune response or MHC class I receptor activity (Suppl. Tables 2 and 3). Signature R1 is particularly interesting, as it shows the highest specificity to the most early stages HSC and MPP, but still appears, though to a lesser degree, in the more differentiated progenitor stages as LMPP, GMP and CMP. We interpret this signature as a stemness signature, which fades away as differentiation progresses and which is absent in the most differentiated cell types. Gene ontology terms associated with this signature were e.g. mesenchymal cell differentiation involvement, cell fate specification or cGMP binding. Genes listed in the MSigDB [29,30] signature *EPPERT_HSC_R* [31], which are up-regulated in HSC enriched populations as compared to committed progenitors and mature cells, are significantly enriched among the genes highly expressed in this signature (p = 2.154 * 10^−14^ as tested with `egsea.ora()` from the R package *EGSEA* [32]). As such, exposures to the signatures reflect developmental trajectories and similarities between the cell populations.

**Figure 2:**
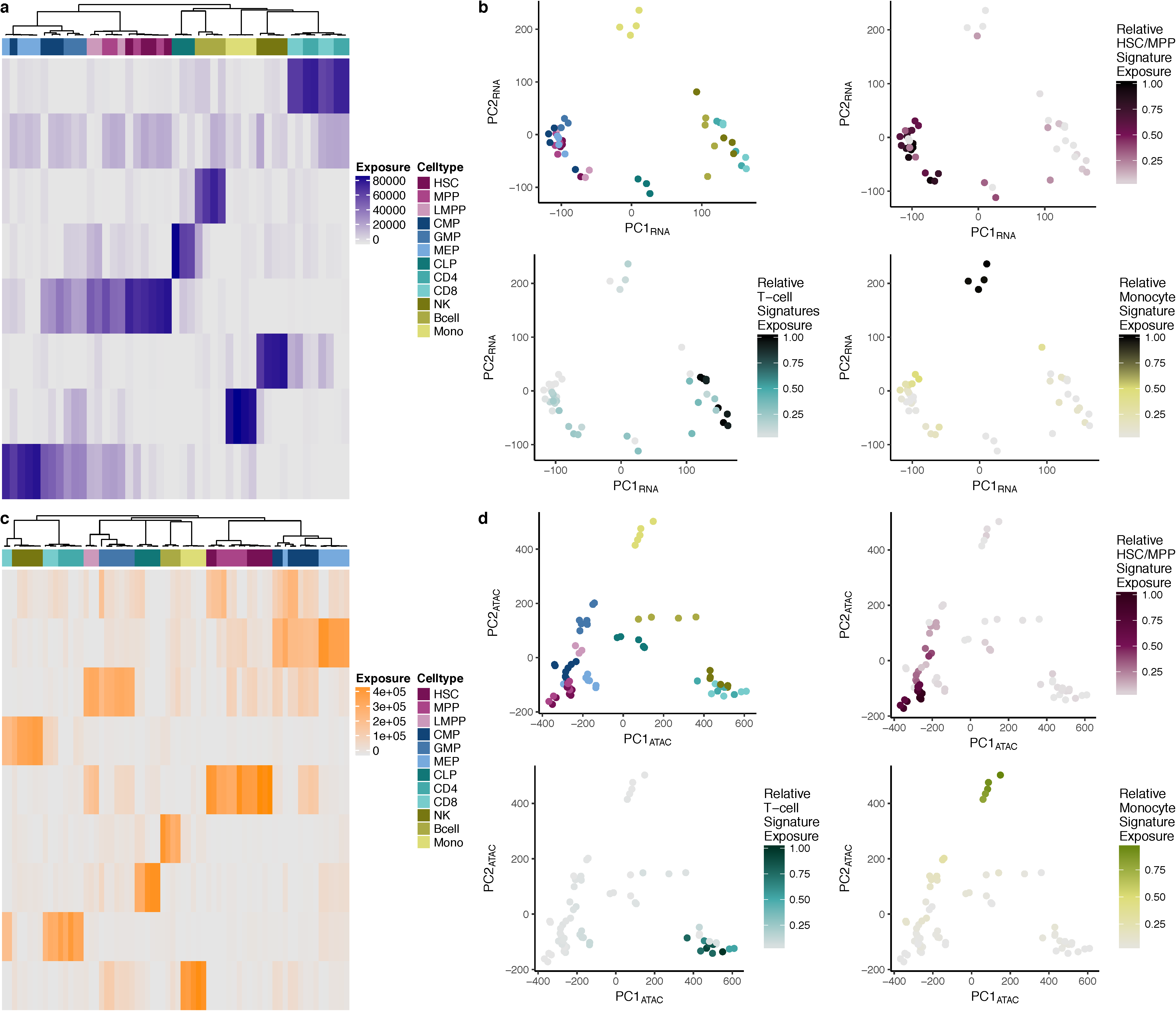
Cell type-resolution dimensionality reduction of RNA- and ATAC-seq datasets by NMF. (a) Heatmap of the matrix H_RNA-seq_ obtained from NMF of an RNA-seq count table from hematopoietic cells. (b) Scatterplot of the first two principal components of the same RNA-seq dataset. Different RNA-seq signatures were mapped onto the PCA visualization to highlight exposure dynamics. (c) Heatmap of the matrix H_ATAC-seq_ obtained from NMF of an ATAC-seq consensus peak count table from hematopoietic cells. (d) Scatterplot of the first two principal components of same ATAC-seq dataset. Different ATAC-seq signatures were mapped onto the PCA visualization to highlight exposure dynamics.

In both decompositions, we found one signature which captures only features (genes or regions) with low specificity, such as e.g. housekeeping genes. Concordantly, gene ontology enrichment analysis yielded housekeeping activities including mitochondrion organization, organelle organization, intracellular transport, RNA processing or DNA repair (Suppl. Tables 2 and 3).

We compared the NMF decomposition to other dimension-reduction approaches, such as PCA. Applying PCA to the same input matrices that were used for NMF, we observed a separation of the cell types according to the first and second principal components (Fig 2b). In order to interpret PCs, we colored the samples according to their exposures to different NMF signatures, the HSC/MPP, the T-cell and the monocyte signatures (Fig 2b). While the separation between myeloid and lymphoid populations appeared clearly along PC1, there was no fine-grained separation of the cell types; for example, the NK and T-cells remained closely clustered together, while they were characterized by distinct signatures in the NMF decomposition. Even when we included a number of PCs comparable to the number signatures, we still could not recover a one-to-one correspondence between PCs and cell populations. For example, there was no PC in RNA-seq or ATAC-seq which distinguished NK cells from the other populations (Supplemental Figure S1). Hence, NMF-derived signatures showed a higher discriminatory power between biological groups (cell types). In addition, signatures followed a hierarchy of prominence in the underlying data. To display this, we combined NMF with a Sankey diagram or riverplot visualization which displays this hierarchy between signatures extracted at different factorization ranks (Supplementary Figures S3 and S4 and Methods section). Signatures extracted at all iterated factorization ranks were compared by non-negative least squares and ordered in a tree-like representation which minimizes edge weights. Signatures extracted at the optimal factorization rank were defined as reference signatures. Signatures extracted at all other factorization ranks were compared to these reference signatures, inheriting their labels and colour codes. At the top of the riverplot for RNA-seq (at rank 2, corresponding to 2 signatures), one myeloid (R3) and one T-cell signature (R5) were found. At rank 3, the next signature which was extracted was the monocyte signature (R8), followed by the B-cell signature (R6) at rank 4, the stemness signature (R1) at rank 5, the NK cell signature (R7) at rank 7 and the diffuse signature (R8) at rank 8.

The ATAC-seq signatures clearly distinguished the various cell populations, achieving a fine-grained description of the samples (Fig. 2c). Compared to the RNA-seq analysis, we found a very similar signature decomposition, with one additional signature. Interestingly, this additional signature showed a high exposure both in LMPP and in GMP, highlighting the relation between these two cell populations.

Visualizing ChIP-seq data from ENCODE [4] and ROADMAP [5] showed that the regions defined by the single-signature features from our ATAC-seq signatures were characterized by enrichment of H3K27ac and H3K4me1 and depletion of H3K27me3, H3K4me3 and H3K9me3 in the differentiated cell types (Fig. 3 for monocytes, Suppl. Fig. S5 for other cell types: first page for CD4-positive T-cells, second page for B-cells, third page for NK cells, fourth page for CD8-positive T-cells). The regions identified in these cell types thus showed epigenetic enhancer hallmarks [33].

**Figure 3:**
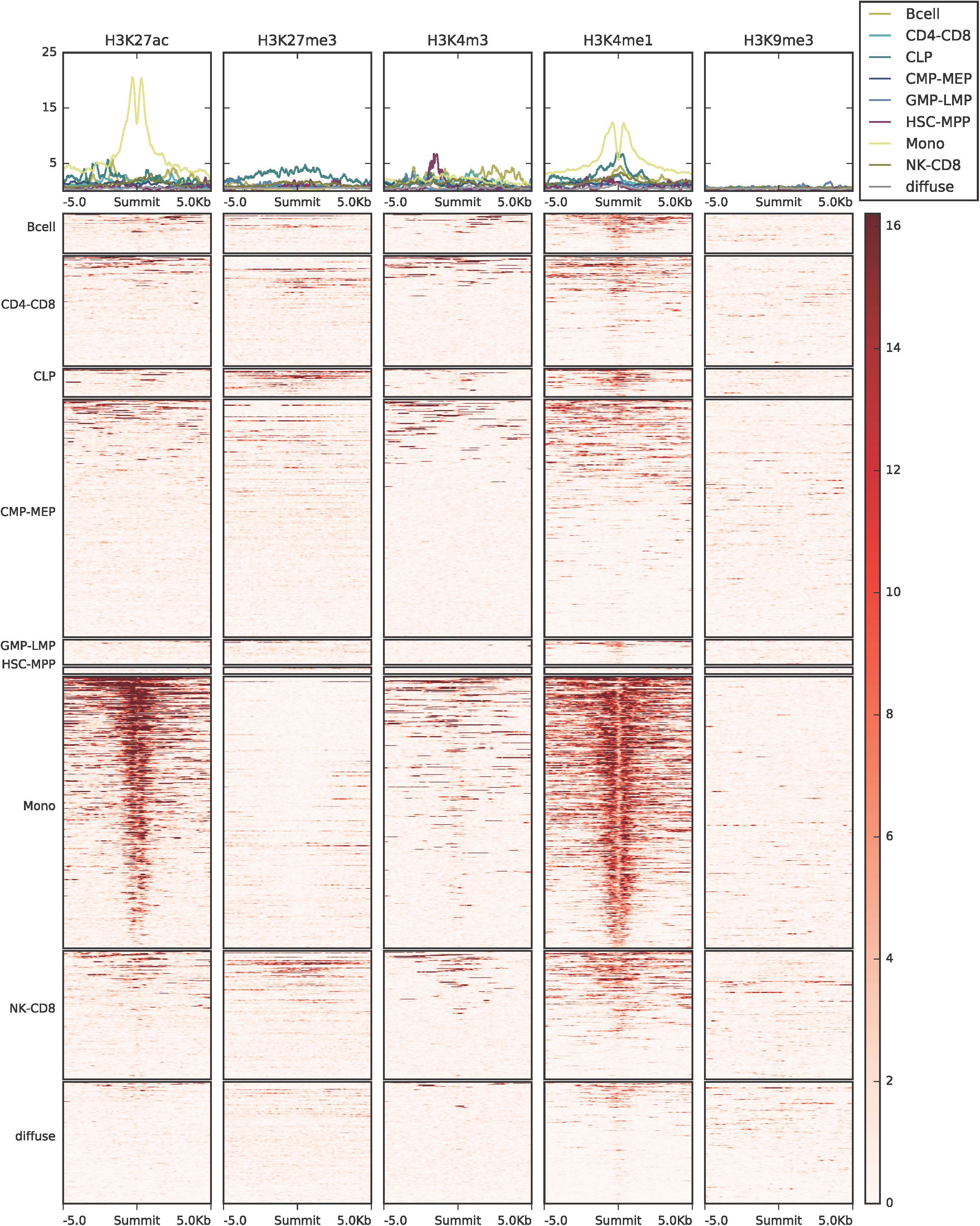
Roadmap ChIP-seq data for monocytes in the single-signature ATAC-seq regions. ChIP-seq signal, as fold-change over control, was plotted for five different histone PTMs, including H3K27ac, H3K27me3, H3K4me3, H3K4me1 and H3K9me3. *Upper panels:* Profile plots showing the ChIP-seq signal in monocytes averaged over single-signature features, i.e. specific regions, for each ATAC-seq signature. *Lower panels:* Heatmaps illustrating the spatial distribution of the ChIP-seq signal +/- 5kb around the peak summit, where each row corresponds to a region and each panel to an ATAC-seq signature.

Given the comparable number of signatures in both datasets, we anticipated that there is a profound relationship between these signatures. In order to understand the origin of this almost one-to-one matching, we extracted the features (genes and genomic regions) corresponding to the RNA-seq and ATAC-seq signatures. We plotted the signal of the single-signature features across all datasets (gene expression for the specific genes and ATAC-seq signal for the specific regions, Fig 4a,b). Apart from the additional LMPP/GMP ATAC-seq signature A2, we again saw a clear overall correspondence between the signatures. However, we also observed interesting differences. For example, the RNA-seq signature R3 showed a specific expression only in MEP, while the corresponding ATAC-seq signature A3 showed a high signal in MEPs and to a lesser extend in CMPs. This is interesting as CMPs are upstream of MEPs, and these regions could correspond to regulatory elements which are primed at the CMP stage and get activated at the MEP stage. To test this hypothesis with publicly available data, we plotted the ChIP-seq signals H3K27me3, H3K4me1, H3K4me3 and H3K9me3 in CMPs across the single-signature features of the different ATAC-seq signatures (Suppl. Fig. S6, first page). Among generally weak signals for the CMP signature A3, there was a slight peak in H3K4me1. In the absence of publicly available human H3K27ac data for CMPs and MEPs, a complementary analysis was performed: we investigated bidirectional transcription with publicly available FANTOM5 data [6,34,35] (Suppl. Fig. S6, second page). In contrast to the differentiated cell types (Suppl. Fig. S7, first page for monocytes, second page for CD4-psotive T-cells, third page for B-cells, fourth page for NK cells, fifth page for CD8-positive T-cells), the single-signature feature regions of the CMP signature A3 showed no clear pattern of bidirectional transcription. Taken together with solely H3K4me1 in ChIP-seq, this is well compatible with primed enhancers at the CMP stage.

**Figure 4:**
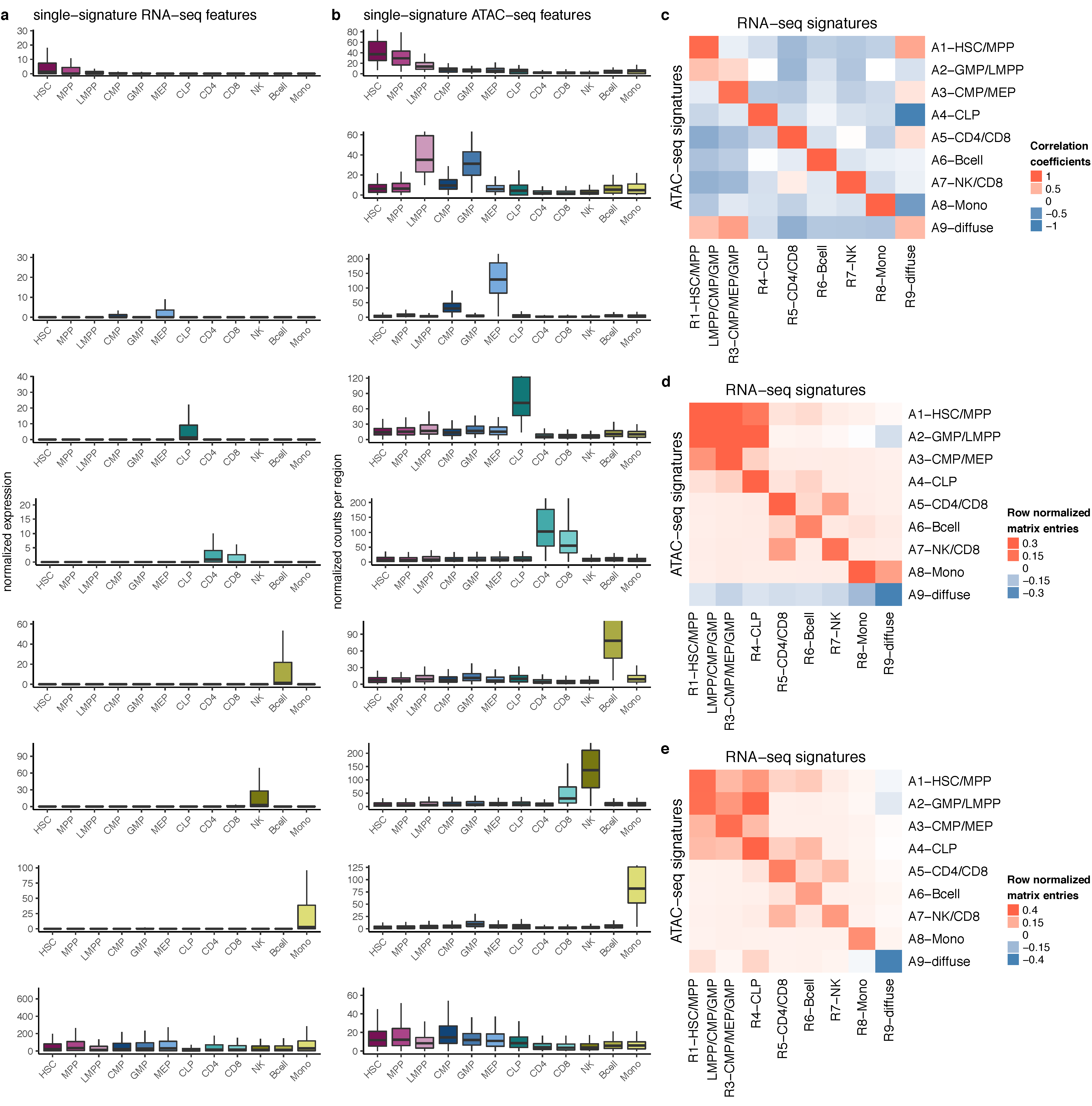
Feature extraction and association after NMF. (a) Boxplots across cell types of signature-specific genes extracted from the matrix W_RNA-seq_. (b) Boxplots across cell types of signature-specific open chromatin regions extracted from the matrix W_ATAC-seq_. (c) Heatmap illustrating pairwise correlations of the rows of the matrices *H* from RNA-seq and ATAC-seq factorizations. (d) Heatmap illustrating pairwise comparisons of the identified signatures, i.e. the columns of the matrices *W* from RNA-seq and ATAC-seq factorizations. As the columns of the matrices *W* of the two omics layers are vectors of different lengths, one omics layer has to be transformed. Such a transformation was obtained using pairwise genomic distances between the features of the two different omics layers (genes for RNA-seq and regions for ATAC-seq). (e) Heatmap illustrating pairwise comparisons of the identified signatures similar to (d), but using only the promoters, i.e. a subset of the ATAC-seq features.

To support the qualitative matching between the signatures and to perform a multi-omics integration, correlations between the exposures to the RNA-seq and ATAC-seq signatures (i.e. H_RNA-seq_ and H_ATAC-seq_; Fig 4c) and comparisons of the signature matrices (i.e. W_RNA-seq_ and W_ATAC-seq_; Fig 4d and 4e, where Fig 4d was generated using all ATAC single-signature features and Fig 4e was generated using only the promoters) were used. The largest correlations of exposures and signatures themselves were observed between corresponding pairs of RNA/ATAC signatures.

To study the degeneracy, i.e. the distribution of redundancy among the features of the signatures identified for RNA-seq and ATAC-seq, we counted single-signature features and multi-signature features (resolving dual-signature features, triple-signature features etc. up to the respective maximum levels of degeneracy, i.e. eight for RNA-seq and nine for ATAC-seq). These counts are shown as barplots in Figure 5b. We observed that in both omics layers, the relative majority of features showed the highest possible level of degeneracy, i.e. they were not specific (high bar in the right of Figure 5b). In order to highlight gross differences between different types of regulatory elements, we labelled ATAC-seq peaks as promoters if they were located within 500 bp of the start of an exon. All other ATAC-seq peaks were labelled as enhancers. Using this definition, we found that a very high fraction of promoters were open in all cell types, whereas the fraction of enhancers which were open in all cell types was lower. When focusing on the remainder of the distributions beyond the strong peak at the right end of the spectrum, further striking differences were observed: whereas genes in RNA-seq showed decreasing counts with increasing levels of degeneracy, ATAC-seq peaks followed a bell-shaped curve and reached their maximum count in the middle of the spectrum. This may highlight a general principle in the hematopoietic lineage: regulation of cell type specific gene expression occurs mainly via combination of regulatory elements which taken individually are neither exclusive nor specific to the cell types [8].

**Figure 5:**
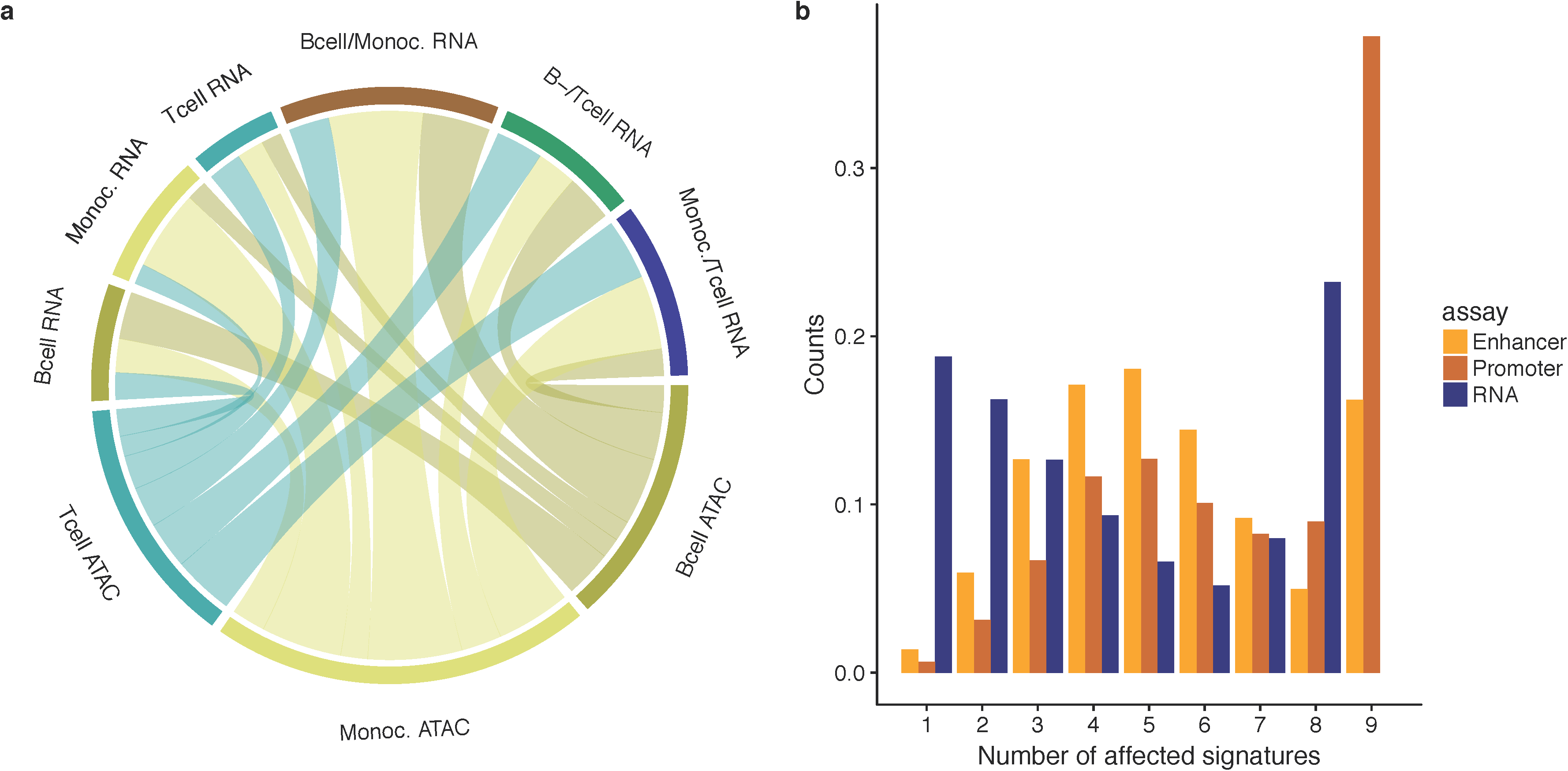
Association of signature-specific ATAC-seq regions with gene expression signature-memberships. (a) Chord diagram of proportions of interactions mapping from signature-specific ATAC-seq regions to genes and their respective signature membership for T-cells, B-cells and monocytes. (b) Barplot showing fractions of features (genes for RNA-seq, blue, and regions for ATAC-seq, dark orange for promoters and light orange for enhancers) specific for combinations of signatures. The x-axis displays the level of degeneracy, i.e. 1 corresponds to single-signature features, 2 to dual-signature features, 3 to triple-signature features etc. up to the respective maximum levels of degeneracy, i.e. eight for RNA-seq and nine for ATAC-seq.

It is well known that regulatory elements do not always target the closest gene, but might interact with genes located more distally. To get a more accurate view of the regulatory interactions, we took advantage of a recently published dataset of chromatin interactions obtained through promoter-capture Hi-C (PCHi-C) in 17 sorted primary blood cell populations, three of which matched cell types of this analysis [36]: B-cells, T-cells and monocytes, to which we restrict the analysis in the following. Starting from the single-signature features for the three corresponding ATAC-seq signatures we mapped these elements using PCHi-C to their respective target genes and recorded which RNA-seq signatures these genes were specific to. As expected, a large portion of single-signature ATAC regions mapped to genes belonging to the corresponding RNA-seq signature (Fig 5a). Conversely, starting from the genes which are specific to the RNA-seq signatures involving T-, B-cells and monocytes, the majority of them interacted with ATAC-seq regions which were specific to the corresponding RNA-seq signature. Two examples of such interactions are shown in Fig. 6.

**Figure 6:**
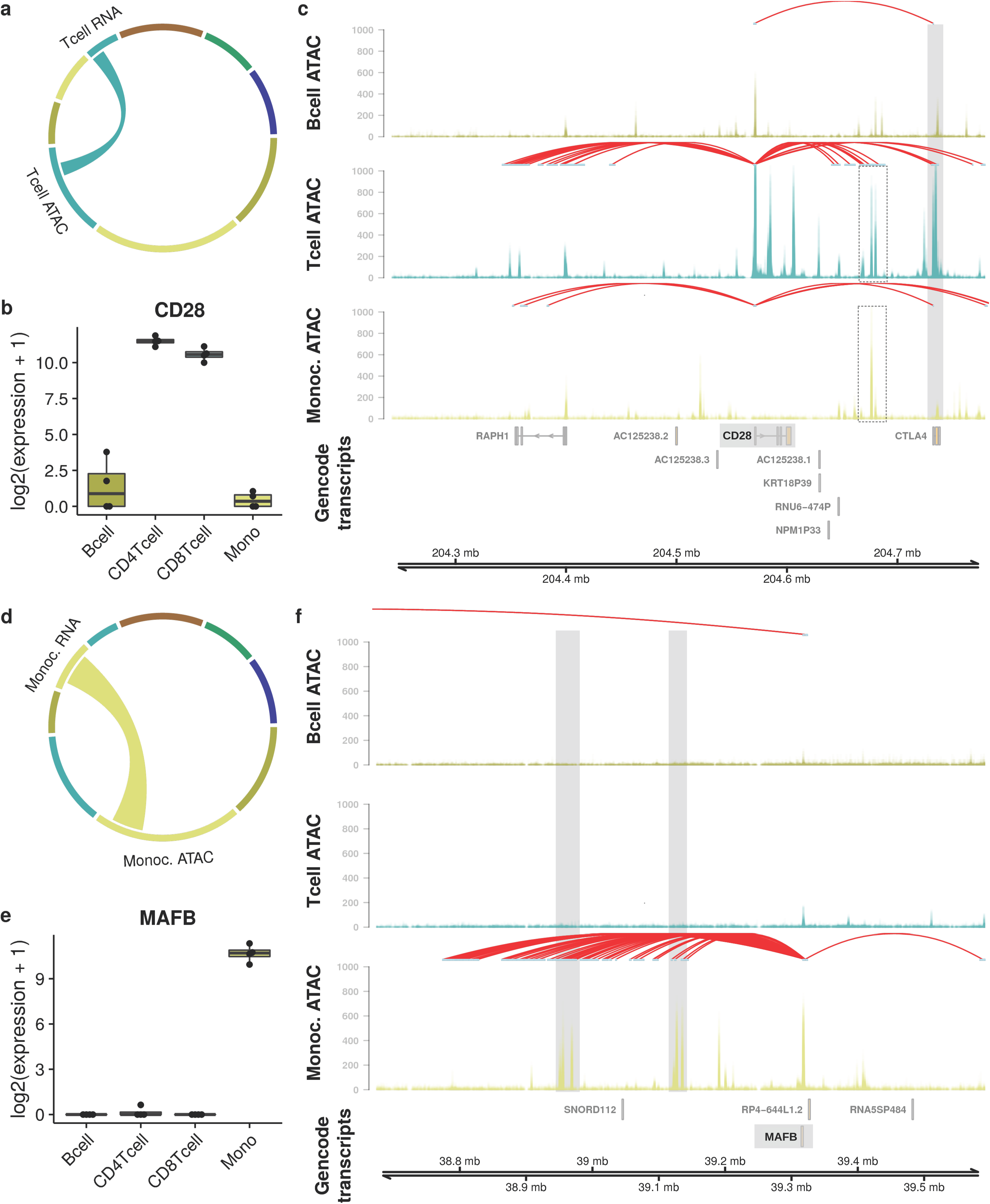
Extraction of cell type-specific open chromatin regions that interact with uniquely expressed genes. (a) Chord diagram indicating the association of T-cell-specific open chromatin regions with genes uniquely expressed in T-cells. (b) Boxplots of log2 transformed library size normalized expression values of CD28 across B-cells, CD4-positive T-cells, CD8-positive T-cells and monocytes. (c) Visualization of the genomic region around CD28 and differential, interacting open chromatin regions. (d) Chord diagram indicating the association of monocyte specific open chromatin regions with genes uniquely expressed in monocytes. (e) Boxplots of log2 transformed library size normalized expression values of MAFB across B-cells, CD4-positive T-cells, CD8-positive T-cells and monocytes. (c) Visualization of the genomic region around MAFB and differentially interacting open chromatin regions.

CD28 had a clear T-cell-specific expression pattern (Fig. 6b). Among the three cell types of interest, this gene had the largest number of interactions to regulatory regions in T-cells, and only one common interaction across all cell types, mapping to a genomic region in the promoter of the gene CTLA4. This region appeared to be widely open in T-cells (middle panel of Fig. 6c), with a very reduced ATAC-seq signal in B-cells and monocytes. Hence the promoters of CD28 and CTLA4 interacted in several cell-types, but the regulatory region was open only in T-cells. The cell-type specificity was thus provided by the openness of chromatin only. This locus also exhibited another mechanism, in which the specificity was provided solely by chromatin conformation: another ATAC-seq peak (dashed rectangle in Fig. 6c), located in the intergenic region between CTLA4 and CD28, showed openness in T-cells and monocytes. Yet only T-cells exhibited an interaction with this potential enhancer, whereas monocytes did not.

The gene MAFB [37] was a single-signature feature for the monocyte RNA-seq signature. Exclusively in monocytes, this gene was subject to PCHi-C interactions with several loci, two of which (highlighted in grey in Fig 6f) belonged to the monocyte-specific ATAC-seq signature.

However, beside these interactions showing a positive regulation, a significant proportion of ATAC-seq regions specific to a particular cell type mapped to genes that were specific for another cell type. Roughly 10% of the monocyte specific ATAC-seq regions mapped to genes that are B-cell-specific (Fig. 5b). Two examples of such hybrid interactions are shown in Fig. 7.

**Figure 7:**
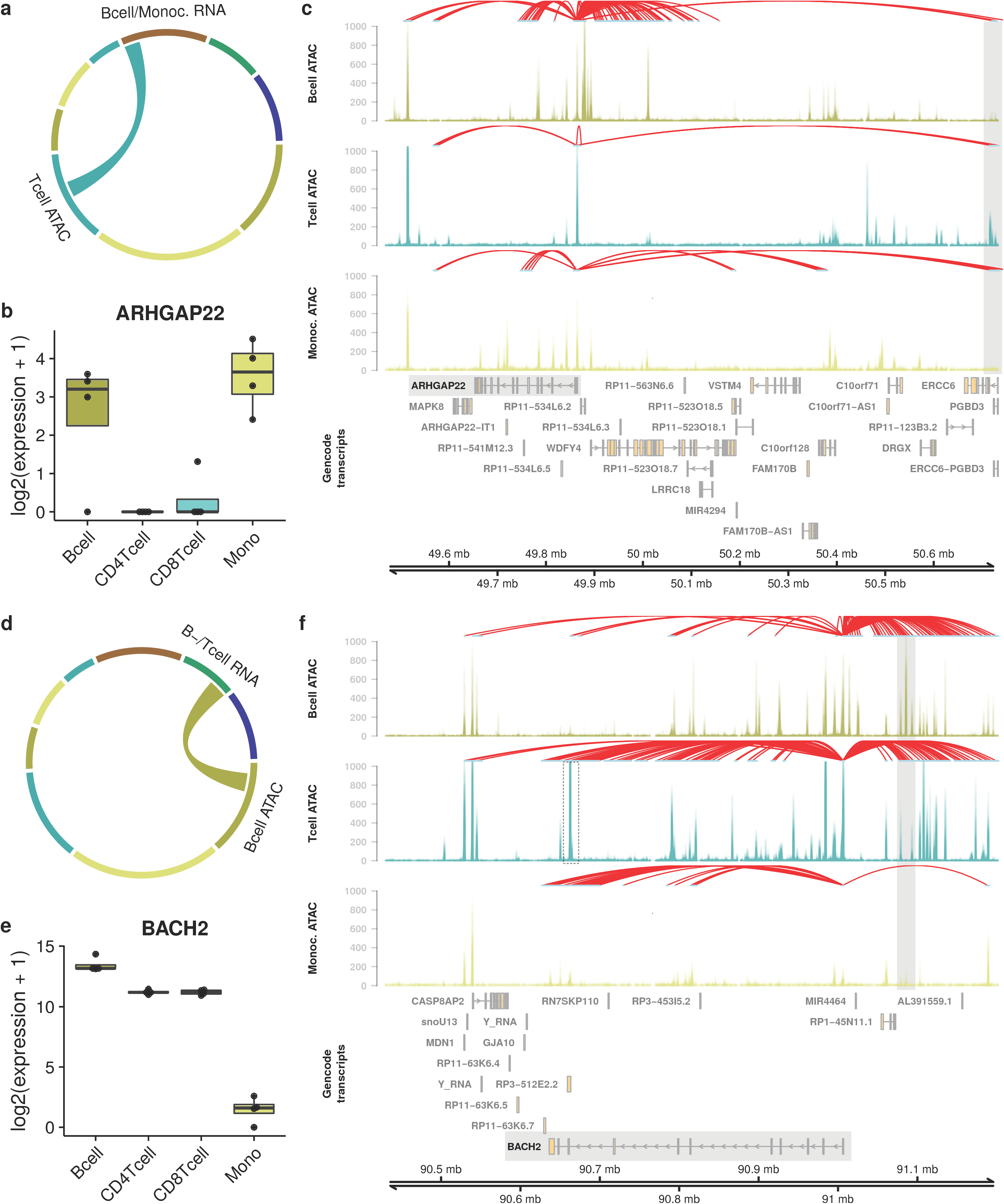
Extraction of cell type-specific open chromatin regions that interact with genes expressed in multiple (other) cell types. (a,d) Chord diagrams indicating which associations of cell type-specific open chromatin regions with which genes are examined. (b,e) Boxplots of log2 transformed library size normalized expression values across B-cells, CD4-positive T-cells, CD8-postive T-cells and monocytes of the genes of interest. (c,f) Visualization of the genomic region around the genes of interest and differentially interacting open chromatin regions.

The gene ARHGAP22, a Rho-GTPase activating protein, was specifically expressed in B-cells and monocytes, but silent in T-cells. One specific locus in the promoter of the ERCC6 gene (highlighted in grey, Fig 7c), which mapped to ARHGAP22 in PCHi-C, showed a striking T-cell specific ATAC-seq signal. This inverted expression/chromatin pattern suggests that this specific region is a silencer element in T-cells. Indeed, ENCODE ChIP-seq in GM12878 showed a peak for RUNX3 at this position, which has been described previously as a repressor of CD4 in T-cell lineage decisions [38]. Of note, ARHGAP22 is included in a predictive 17 gene score for AML risk estimation [39] and has been linked to regulation of tumor cell mobility [40].

The gene BACH2 was expressed in both B-cells and T-cells (Fig. 7e), and displayed a complex array of interactions in these two cell types. Interestingly, many of these connected regions showed a pattern which was specific of either cell type; the highlighted region (Fig. 7f) was strongly and specifically open in B-cells but closed in T-cells. Another region (close to the 5’ end of the BACH2 transcript, highlighted by a dashed rectangle in Fig. 7f) showed an opposite pattern of strong T-cell signal, and no signal in either B-cells or monocytes. This indicates that the transcriptional regulation occured here via very distinct regulatory programs.

## Discussion

We presented a new workflow implemented into an R package called Bratwurst (available at https://github.com/wurst-theke/bratwurst), which implements a novel NMF workflow for genomic datasets. Through wrapper functions for NMF solvers implemented in CUDA, it makes the highly parallel architecture of GPUs and efficient matrix operations available to decompositions of large matrices in R. One major advantage of NMF is that both matrices issued from the factorization, the *signatures matrix W* and the *exposure matrix H*, can be used for data analysis and interpretation. Notably, we use the *exposure matrix H* for a comparison with labels of the samples for recovery of predefined subgroups and thus extraction of subgroup-specific signatures.

The extracted signatures correspond to subgroup-specific patterns. Here, we applied the functionality provided by the Bratwurst package to published RNA-seq and ATAC-seq data obtained from multiple hematopoietic cell types [22]. Signatures extracted from RNA-seq correspond to transcriptional programs and those extracted from ATAC-seq data correspond to regulatory programs. Relating signatures extracted from the two different omics layers corresponds to relating transcriptional to regulatory programs well beyond relating single genes to single regulatory elements. The functionality in Bratwurst is in no way limited to these two data types, and can be applied to all high-dimensional data. Additional information, like e.g. chromatin conformation data, may be used to enhance the relationship between the signatures extracted from the different omics layers. Thus, Bratwurst is also a powerful tool for multi-omics integration.

Beyond matrix factorization by NMF and multi-omics comparison, the Bratwurst package provides a new feature selection strategy. It extracts features (i.e. genes in the case of RNA-seq and regions in the case of ATAC-seq) which are specific for single signatures (single-signature features) or for groups of signatures (multi-signature features). This strategy is non-parametric and does not depend on user choices for cutoffs or assumptions of any kind. Extracted features may be used for different purposes in a downstream analysis, including but not limited to focused analyses of single genes and their regulome or enrichment analyses.

The design of the feature selection strategy allows to tune specificity to a user-defined level: single-signature features being most specific, and specificity gradually decreasing with increasingly broad combinations of signatures. The specificity of the whole method scales with increasing factorization rank. Therefore, transparent and reproducible strategies to determine the optimal factorization rank, as implemented in the workflow presented here, are mandatory.

Inclusion of the Sankey diagram or riverplot visualization into the downstream analysis after NMF provides additional information which would otherwise not be available: signatures at different factorization ranks are compared and ordered by their degree of similarity, providing a tree structure showing how additional signatures emerge at increasing factorization ranks. In the application to different cells of hematopoiesis, the above strategy yields new insights into the interplay between open chromatin, chromatin conformation and gene expression. Considering that subtle differences in transcript abundance might be required to establish a certain cellular phenotype, observations as the regulatory landscape of BACH2 (Figure 7d,e,f) indicate that cells make use of such highly complex compositions of regulatory elements to fine tune target gene expression. Different examples highlighted that cell type-specific gene expression can be regulated via cell type-specific chromatin interactions, cell type-specific chromatin openness or combinations thereof. Different cell types use different enhancers to trigger expression of the same gene. Similar transcriptional states are achieved by distinct regulatory programs (Figure 7). The majority of the enhancers which constitute these regulatory programs are not specific to single cell types but are conserved across several cell types instead (Figure 5b). The programs, however, which are combinations of these less specific regulatory elements, are very specific for the different cell types [41]. Additionally, a few cell type-specific regulators are present. The type of analysis presented in this work enables an in depth dissection of this plethora of interacting mechanisms.

The capacities of a package like Bratwurst may become increasingly valuable with the increasing amounts of produced and available data from different omics technologies. Single-cell technologies are revolutionizing the understanding of complex and heterogeneous biological systems, notably in upcoming consortia efforts such as the human cell atlas [42]. Bratwurst may be a suitable tool to extract and relate regulatory and/or transcriptional programs in single-cell data analogously to the bulk data analyzed in this manuscript. As NMF algorithms have recently been successfully applied to single-cell RNA-seq data in order to obtain clusters or subgroups [43,44], our feature extraction methodology could be applied to such factorizations and map features to subgroups. In addition the ability of our method to integrate across multiple omics layers could also benefit the analysis of upcoming datasets with multi-omics measurements in single cells [45–47].

## Conclusion

The Bratwurst software provides functionalities to identify patterns in different types of omics data with NMF, relate the identified patterns to subgroups, select features highly specific to the different patterns and integrate the patterns extracted from the different omics layers. Portation of NMF to GPUs takes advantage of massive parallelization and enables deconvolution of large matrices with NMF. We applied this software to RNA-seq and ATAC-seq data from hematopoietic cells and found distinct cell type-specific and intimately linked regulatory and transcriptional programs. Our software can be applied to any omics data type and is a powerful tool for multi-omics integration.

## Methods

As outlined in the workflow section of the results part of this manuscript, Bratwurst (https://github.com/wurst-theke/bratwurst) provides a multi-step workflow. We herewith provide complementary and detailed descriptions of the different steps and describe choices made for the analysis of RNA-seq data and ATAC-seq data of hematopoietic cells from the Corces et al. dataset. Furthermore, Bratwurst introduces a new S4 class, `nmfExperiment`, which inherits from `SummarizedExperiment` [48].

- **Step 1: Preprocessing**.
  - (a) Preprocessing of the RNA-seq data: after download from ftp://ftp.ncbi.nlm.nih.gov/geo/series/GSE74nnn/GSE74246/suppl/GSE74246%5FRNAseq%5FAll%5FCounts.txt.gz, HSC, MPP, LMPP, CMP, GMP, MEP, CLP, CD4-positive T-cell, CD8-positive T-cell, NK cell, B-cell, and monocyte samples were chosen for analysis. In addition, the sample X5852.GMP was removed, as it had much smaller library size than other GMP samples and was considered as an outlier in this respect. Rows with only zero entries were removed, and a rescaling with the function `estimateSizeFactorsForMatrix()` from the R package DESeq2 [23] was performed. The data were log_2_(x + 1)-transformed and finally stored in the object `rna.nmf.exp` of class `nmfExperiment` (available at https://doi.org/10.5281/zenodo.800049).
  - (b) Preprocessing of the ATAC-seq data: after download from ftp://ftp.ncbi.nlm.nih.gov/geo/series/GSE74nnn/GSE74912/suppl/GSE74912%5FATACseq%5FAll%5FCounts.txt.gz, samples of the same cell types as for RNA-seq were chosen for analysis and preprocessing was performed analogously. Rows with rowSums < 2000 were removed, and sample X6792.7A was removed due to low coverage. In analogy with preprocessing of RNA-seq data, a rescaling with the function `estimateSizeFactorsForMatrix()` from the R package DESeq2 [23] and a subsequent log_2_(x + 1) transformation were performed. Data from the ATAC-seq analysis is stored in the object `atac.nmf.exp` of class `nmfExperiment` (available at https://doi.org/10.5281/zenodo.800049).
- **Step 2: NMF**. As illustrated in the center of Figure 1, the task of NMF algorithms is to factorize one large matrix *V* (of dimensions *n* x *m*) into two smaller matrices W (the *signature matrix* of dimensions *n* x *k*) and *H* (the *exposure matrix* of dimensions *k* x *m*) under the constraint of non-negativity on all entries in both factor matrices *W* and *H* (Figure 1):

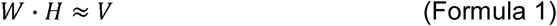 In this work we stick to the following nomenclature: we call the rows of the input matrix *V* and of the signature matrix *W features*. Usually, the columns of *V* (and thus the columns of *H)* correspond to samples. The *exposure matrix H* represents contributions of the *k* identified signatures to the *n* different input samples and may be used for a soft clustering, whereas the *signature matrix W* captures the contributions of the *m* different features to the *k* signatures. In order to achieve a reduction in complexity and dimensionality, the *factorization rank k*, which is a free parameter, has to be chosen s.t. *k* < *n* and *k* < *m*. The task of NMF algorithms may be reformulated as an optimization problem in which the residuals are to be minimized: minimize∥ *V*− *W · H* ∥ under the above-mentioned constraint of non-negativity. In order to solve this optimization task, all NMF algorithms share an internal two-substep procedure: (i) initialization and (ii) iteration over update equations. The different algorithms in the NMF family have different implementations of these internal substeps. In principle, any of these algorithms may be called from Bratwurst. If no GPU is available, Bratwurst by default uses an NMF implementation which is executed on the CPU (R package NMF [25]). If a GPU is available, an NMF implementation taking advantage of this highly parallel architecture for efficient matrix operations can be used by Bratwurst, e.g. NMF_GPU [24](function `runNmfGpu()`). In the following, we present the details of a new implementation of an NMF-solver using the python modules PyCUDA (https://pypi.python.org/pypi/pycuda) and cudamat (https://github.com/cudamat) as interface to the NVIDIA CUDA 8 toolkit. This solver is called by a wrapper function from the R package Bratwurst (function `runNmfGpuPyCuda()`):
  - (i) The initialization is performed by filling initial instances of the matrices *W* and *H* with random numbers.
  - (ii) Iteration until convergence over update equations as used in [9].
  One run of iterations over update equations after choosing one set of initial conditions yields one solution (if leading to convergence). In order to sample the available parameter space, however, it is necessary to choose different initial conditions and iterate update equations until convergence starting from these. Among all tested initial conditions, the one with the lowest residual is chosen. Therefore, the steps (i) and (ii) described above are wrapped into an iteration over initial conditions. This gives NMF a stochastic character. As the factorization rank *k* is a free parameter for any NMF method, Bratwurst determines the optimal factorization rank by iterating over different factorization ranks, evaluating quality metrics for any one of them and selecting the rank which complies best with the quality metrics [26]. In this work, we chose to:
  - minimize the Frobenius norm of the residuals (function `computeFrobErrorStats()`)
  - maximize the cophenetic correlation coefficient [10] (function `computeCopheneticCoeff()`) and
  - minimize the Amari-type distance [14] (function `computeAmariDistances()`).
  The function `proposeOptK()` gives a suggestion for the optimal factorization rank. Therefore, in an analysis with Bratwurst, we follow a threefold nested iterative procedure:
  - (1) iteration over factorization ranks
  - (2) iteration over initial conditions
  - (3) iteration over update equations
  The class `nmfExperiment` provides slots to store the factorized matrices *W* and *H* from every single factorization. It is important to keep this data in order to be able to compute the quality metrics described above. The matrices are stored in a named list of lists (the upper level is named by factorization rank, the lower level by the iterating index over initial conditions), for which the accessor functions `HMatrix()`, `HMatrixList()`, `WMatrix()` and `WMatrixList()` are provided. For the analyses of both the RNA-seq (a) and ATAC-seq (b) data from Corces et al. [22], update equations were iterated 2 · 10^4^ times unless convergence was reached earlier (checked every 10 iterations). 200 iterations over initial conditions were performed. Iterations over factorization ranks covered the range 2 - 14. Signatures extracted at different factorization ranks can be visualized in a Sankey diagram or riverplot [27]. In this two-dimensional tree representation (function `generateRiverplot()`), the nodes correspond to the extracted signatures and the edge weights are determined by computing non-negative least squares decompositions of the signatures at factorization rank *r* with the signatures at the next lower factorization rank *r - 1*. The y-coordinate encodes factorization rank (increasing from top to bottom), whereas the x-coordinate denotes the signatures extracted at one fixed factorization rank. The nodes, corresponding to the signatures, are ordered according to similarity across factorization ranks, leading to thick edges being short and thin edges being long. A matrix factorization of type ☐ · ☐, ≈ ☐ is invariant under scalar transformations with a constant factor *c*: 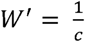 and *H*′ = *c* · *H*. This ambiguity can be resolved by imposing the requirement that *W* is column-normalized. In order to keep the data stored in an object of class `nmfExperiment` consistent, it is important to perform the column normalization of *W* and corresponding rescaling of *H* on all matrices stored in the object (originating from the different iterations) simultaneously. This is done by the function `normalizeW()`. The normalized versions of (a) the RNA-seq data and (b) the ATAC-seq data are stored in the objects `norm.rna.nmf.exp` and `norm.atac.nmf.exp`, respectively (also available for download at https://doi.org/10.5281/zenodo.800049)
- **Step 3: Mapping signatures to labelled groups of samples**. This step is described in detail in the workflow section of the results part of this manuscript. After having mapped signatures to subgroups of samples, the riverplot generated in the previous step can be updated. To this end, the signatures extracted for the optimal factorization rank are defined to be reference signatures. A colour coding of these reference signatures may be inherited from sample annotations. All signatures in the riverplot, i.e. signatures extracted at all iterated factorization ranks, are then labelled and coloured according to the most similar reference signature, where similarity is computed by cosine similarity (function `attributeComparisonSignatures()`).
- **Step 4: Feature selection**. This step is described in detail in the workflow section of the results part of this manuscript. The extracted lists of single-signature features and multi-signature features are stored in specific slots of `norm.rna.nmf.exp` and `norm.atac.nmf.exp`, respectively (available for download at https://doi.org/10.5281/zenodo.800049). Enrichment analyses can be applied to the feature lists extracted as described above.

The analysis (Steps 1-4) was run individually for RNA-seq and ATAC-seq data. In order to relate these two layers of data, pairwise comparisons between the exposures (rows of the matrices H_RNA-seq_ and H_ATAC-seq_) and the signatures (columns of the matrices W_RNA-seq_ and W_ATAC-seq_) were performed. Exposures of the two different omics layers were compared by pairwise correlations of the rows of the exposure matrices:

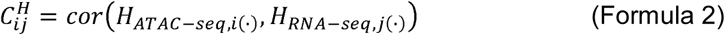

where *H*_*ATAC-Seq,i*(·)_ is the i^th^ row of the matrix *H*_*ATAC*-Seq_and *H*_*RNA-Seq,j*(·)_ is the j^th^ row of the matrix H_*RNA-seq*_.

The signatures, i.e. the columns of the matrices W of the two omics layers, are vectors of different lengths. In order to compare these by pairwise correlation, one omics layer has to be transformed by a suitable metric M. Such a metric can be obtained by first computing pairwise genomic distances between the features of the two different omics layers (genes for RNA-seq and regions for ATAC-seq):

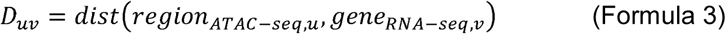

Next, a suitable function of the distances is applied element-wise to this matrix. According to Ouyang et al. [49], this function can be chosen as

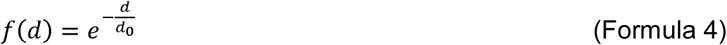

where d_0_ is a characteristic distance evaluated to be 5000 bp in [49]. We can then compute the comparison of the matrices *W* as:

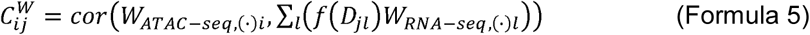

where *w*_*ATAC–seq*,(·)*i*_ is the i^th^ column of *w*_*ATAC*–Seq_and *w*_*RNA-seq*,(·)*i*_ is the j^th^ column of *W_RNA-seq_*. Values in the resulting matrix M can be colour coded in a heatmap.

For three cell types of this analysis (B-cells, T-cells and monocytes), chromatin interaction data obtained through promoter-capture Hi-C [36] was available. After download from http://www.cell.com/cms/attachment/2086554122/2074217047/mmc4.zip, data preprocessing and filtering was performed with the CHiCAGO method [50].

## Acknowledgements

This work was supported by the BMBF-funded Heidelberg Center for Human Bioinformatics (HD-HuB) within the German Network for Bioinformatics Infrastructure (de.NBI) (#031A537A, #031A537C). D.H. acknowledges support by Prof. Dr. M. Lanzer and Dr. C. Denk in the framework of the MD/PhD-program of Heidelberg University.

## Author contributions

D.H. and S.S. initiated software development, N.K. and C.H. chose the data, N.K. and D.H. performed the analysis, N.K., D.H., S.S., P.R., S.K., C.A. and J.P. developed code for the software package, N.K., D.H., C.H. and M.S. wrote the manuscript, C.H., M.S. and R.E. supervised the project. All authors critically reviewed the manuscript.

---

**Supplementary Figure S1: Heat maps of first nine principal components of RNA- and ATAC-seq datasets.** (a) RNA-seq (b) ATAC-seq, row labels correspond to variance explained by respective principal components.

**Supplementary Figure S2: Quality metrics of RNA- and ATAC-seq dataset NMF with different factorization ranks.** Frobenius errors, cophenetic coefficients and mean Amari distance for (a) RNA-seq and (b) ATAC-seq factorizations. The chosen optimal factorization ranks are highlighted by grey rectangles.

**Supplementary Figure S3: Riverplot for RNA-seq NMF.** Tree-like representation of the signatures extracted from RNA-seq of hematopoietic cells at different factorization ranks. Nodes represent the signatures, the edge strength encodes cosine similarity between signatures linked by the edges. The factorization rank increases from top to bottom. After choosing the optimal factorization rank k_RNA-seq_ = 8 (highlighted by a rectangle), the signatures extracted at this rank are called reference signatures and colours are assigned to them. Then, every signature extracted at every other iterated factorization rank is compared to the reference signatures and coloured by the attributes of the most similar reference signature. Labels of the reference signatures with the suffix “-like” are assigned to the signatures.

**Supplementary Figure S4: Riverplot for ATAC-seq NMF.** Representation analogous to Supplementary Figure S3 of the signatures extracted from ATAC-seq of hematopoietic cells at different factorization ranks. Reference signatures are assigned at the optimal factorization rank k_ATAC-seq_ = 9 (highlighted by a rectangle).

**Supplementary Figure S5: Roadmap ChIP-seq data for differentiated cell types in the single-signature ATAC-seq regions.** First page: CD4-positive T-cells, second page: B-cells, third page: NK cells, fourth page: CD8-positive T-cells. ChIP-seq signal, as fold-change over control, was plotted for five different histone PTMs, including H3K27ac, H3K27me3, H3K4me3, H3K4me1 and H3K9me3, analogous to main Figure 4. *Upper panels:* Profile plots showing the ChIP-seq signal in the respective cell types averaged over single-signature features, i.e. specific regions, for each ATAC-seq signature. *Lower panels:* Heatmaps illustrating the spatial distribution of the ChIP-seq signal +/- 5kb around the peak summit, where each row corresponds to a region and each panel to an ATAC-seq signature.

**Supplementary Figure S6: Combined epigenetic data for CMPs in the single-signature ATAC-seq regions.** First page: ChIP-seq signal, as fold-change over control, was plotted for five different histone PTMs, including H3K27me3, H3K4me3, H3K4me1 and H3K9me3. *Upper panels:* Profile plots showing the ChIP-seq signal in CMPs averaged over single-signature features, i.e. specific regions, for each ATAC-seq signature. *Lower panels:* Heatmaps illustrating the spatial distribution of the ChIP-seq signal +/- 5kb around the peak summit, where each row corresponds to a region and each panel to an ATAC-seq signature. Second page: Metaprofiles of CAGE-seq data from FANTOM5 in CMPs. Transcript per million (TPM) normalized tags were summed up over single-signature regions in the different ATAC-seq signatures A1 - A9. Tags on the minus strand were multiplied by -1 for visualization purposes. Tags on the minus strand are displayed in blue, whereas tags on the plus strand are displayed in red.

**Supplementary Figure S7: Metaprofiles of CAGE-seq data from FANTOM5 in differentiated cell types.** Transcript per million (TPM) normalized tags were summed up over single-signature regions in the different ATAC-seq signatures A1 - A9. Tags on the minus strand were multiplied by -1 for visualization purposes. Tags on the minus strand are displayed in blue, whereas tags on the plus strand are displayed in red. First page: monocytes, second page: CD4-positive T-cells, third page: B-cells, fourth page: NK cells, fifth page: CD8-positive T-cells.

**Supplementary Table T1: Information about the samples used.** Numbers of samples for RNA-seq, ATAC-seq and their overlap.

**Supplementary Table T2: Results of the topGO enrichment for biological processes (gene ontology class *BP*) of the single-signature features from RNA-seq.**

**Supplementary Table T3: Results of the topGO enrichment for molecular functions (gene ontology class *MF*) of the single-signature features from RNA-seq.**

**Supplementary Table T4: Single signature features of the RNA-seq signatures (gene lists).**

**Supplementary Files B1 - B9: Single signature features of the ATAC-seq signatures (BED files).** File B1: ATAC-seq signature *A1-HSC-MPP*; file B2: signature *A2-GMP-LMPP*; file B3: signature *A3-CMP-MEP*; file B4: signature *A4-CLP*; file B5: signature *A5-CD4-CD8*; file B6: signature *A6-Bcell*; file B7: signature *A7-NK*; file B8: signature *A8-Mono*; file B9: signature *A9-diffuse*.

## References

[1] Levine M, Tjian R. Transcription regulation and animal diversity. Nature 2003;424:147–51.

[2] Lenhard B, Sandelin A, Carninci P. Metazoan promoters: emerging characteristics and insights into transcriptional regulation. Nat Rev Genet 2012;13:233–245.

[3] Bulger M, Groudine M. Enhancers: the Abundance and Function of Regulatory Sequences Beyond Promoters. Dev Biol 2010;339:250–7. doi:10.1016/j.ydbio.2009.11.035.Enhancers.

[4] Feingold E, Good P, Guyer M, Kamholz S, Liefer L, Wetterstrand K, et al. The ENCODE (ENCyclopedia Of DNA Elements) Project. Science (80-) 2004;306:636–40. doi:10.1126/science.1105136.

[5] Bernstein BE, Stamatoyannopoulos JA, Costello JF, Ren B, Milosavljevic A, Meissner A, et al. The NIH Roadmap Epigenomics Mapping Consortium. Nat Biotechnol 2010;28:1045–1048.

[6] Andersson R, Gebhard C, Miguel-Escalada I, Hoof I, Bornholdt J, Boyd M, et al. An atlas of active enhancers across human cell types and tissues. Nature 2014;507:455–461.

[7] Stunnenberg HG, The International Human Epigenome Consortium, Hirst P. The International Human Epigenome Consortium: A Blueprint for Scientific Collaboration and Discovery. Cell 2016;167:1145–9.

[8] Lara-Astiaso D, Weiner A, Lorenzo-Vivas E, Zaretsky I, Jaitin DA, Eyal D, et al. Chromatin state dynamics during blood formation. Science (80-) 2014;345:943–9. doi:10.1126/science.1256271.

[9] Lee DD, Seung HS. Learning the parts of objects by non-negative matrix factorization. Nature 1999;401:788–91. doi:10.1038/44565.

[10] Brunet JP, Tamayo P, Golub TR, Mesirov JP. Metagenes and molecular pattern discovery using matrix factorization. Proc Natl Acad Sci U S A 2004;101:4164–9. doi:10.1073/pnas.0308531101.

[11] Devarajan K. Nonnegative matrix factorization: An analytical and interpretive tool in computational biology. PLoS Comput Biol 2008; 4. doi:10.1371/journal.pcbi.1000029.

[12] Alexandrov LB, Nik-Zainal S, Wedge DC, Aparicio S a JR, Behjati S, Biankin A V, et al. Signatures of mutational processes in human cancer. Nature 2013;500:415–21. doi:10.1038/nature12477.

[13] Alexandrov LB, Nik-Zainal S, Wedge DC, Campbell PJ, Stratton MR. Deciphering Signatures of Mutational Processes Operative in Human Cancer. Cell Rep 2013;3: 246–59. doi:10.1016/j.celrep.2012.12.008.

[14] Wu S, Joseph A, Hammonds AS, Celniker SE, Yu B, Frise E. Stability-driven nonnegative matrix factorization to interpret spatial gene expression and build local gene networks. Proc Natl Acad Sci 2016;113:201521171. doi:10.1073/pnas.1521171113.

[15] Moffitt RA, Marayati R, Flate EL, Volmar KE, Loeza SGH, Hoadley KA, et al. Virtual microdissection identifies distinct tumor- and stroma-specific subtypes of pancreatic ductal adenocarcinoma. Nat Genet 2015;47:1168–78. doi:10.1038/ng.3398.

[16] Pal S, Bi Y, MacYszyn L, Showe LC, O’Rourke DM, Davuluri R V. Isoform-level gene signature improves prognostic stratification and accurately classifies glioblastoma subtypes. Nucleic Acids Res 2014; 42. doi:10.1093/nar/gku121.

[17] Qi Q, Zhao Y, Li M, Simon R. Non-negative matrix factorization of gene expression profiles: A plug-in for BRB-ArrayTools. Bioinformatics 2009;25:545–7. doi:10.1093/bioinformatics/btp009.

[18] Frigyesi A, Höglund M. Non-negative matrix factorization for the analysis of complex gene expression data: Identification of clinically relevant tumor subtypes. Cancer Inform 2008;6:275–92.

[19] Carmona-Saez P, Pascual-Marqui RD, Tirado F, Carazo JM, Pascual-Montano A. Biclustering of gene expression data by Non-smooth Non-negative Matrix Factorization. BMC Bioinformatics 2006;7:78. doi:10.1186/1471-2105-7-78.

[20] Gao Y, Church G. Improving molecular cancer class discovery through sparse non-negative matrix factorization. Bioinformatics 2005;21:3970–5. doi:10.1093/bioinformatics/bti653.

[21] Li YE, Xiao M, Shi B, Yang Y-CT, Wang D, Wang F, et al. Identification of high-confidence RNA regulatory elements by combinatorial classification of RNA–protein binding sites. Genome Biol 2017;18:169. doi:10.1186/s13059-017-1298-8.

[22] Corces MR, Buenrostro JD, Wu B, Greenside PG, Chan SM, Koenig JL, et al. Lineage-specific and single-cell chromatin accessibility charts human hematopoiesis and leukemia evolution. Nat Genet 2016;48:1193–203. doi:10.1038/ng.3646.

[23] Love MI, Huber W, Anders S. Moderated estimation of fold change and dispersion for RNA-seq data with DESeq2. Genome Biol 2014;15:550. doi:10.1186/s13059-014-0550-8.

[24] Mejía-Roa E, Tabas-Madrid D, Setoain J, García C, Tirado F, Pascual-Montano A. NMF-mGPU: non-negative matrix factorization on multi-GPU systems. BMC Bioinformatics 2015;16:43. doi:10.1186/s12859-015-0485-4.

[25] Gaujoux R, Seoighe C. A flexible R package for nonnegative matrix factorization. BMC Bioinformatics 2010;11:367. doi:10.1186/1471-2105-11-367.

[26] Amari S, Cichocki A, Yang HH. A New Learning Algorithm for Blind Signal Separation. Adv Neural Inf Process Syst 1996;8:757–63.

[27] Weiner J. riverplot: Sankey or Ribbon Plots 2017.

[28] Alexa A, Rahnenfuhrer J. topGO: Enrichment Analysis for Gene Ontology 2016.

[29] Liberzon A, Subramanian A, Pinchback R, Thorvaldsdóttir H, Tamayo P, Mesirov JP. Molecular signatures database (MSigDB) 3.0. Bioinformatics 2011;27:1739–40. doi:10.1093/bioinformatics/btr260.

[30] Subramanian A, Tamayo P, Mootha VK, Mukherjee S, Ebert BL, Gillette MA, et al. Gene set enrichment analysis: A knowledge-based approach for interpreting genome-wide expression profiles. Proc Natl Acad Sci 2005;102:15545–50. doi:10.1073/pnas.0506580102.

[31] Eppert K, Takenaka K, Lechman ER, Waldron L, Nilsson B, van Galen P, et al. Stem cell gene expression programs influence clinical outcome in human leukemia. Nat Med 2011;17:1086–93. doi:10.1038/nm.2415.

[32] Alhamdoosh M, Ng M, Wilson NJ, Sheridan JM, Huynh H, Wilson MJ, et al. Combining multiple tools outperforms individual methods in gene set enrichment analyses. Bioinformatics 2017;33:414–24. doi:10.1093/bioinformatics/btw623.

[33] Heintzman ND, Stuart RK, Hon G, Fu Y, Ching CW, Hawkins RD, et al. Distinct and predictive chromatin signatures of transcriptional promoters and enhancers in the human genome. Nat Genet 2007;39:311–8. doi:10.1038/ng1966.

[34] Lizio M, Harshbarger J, Shimoji H, Severin J, Kasukawa T, Sahin S, et al. Gateways to the FANTOM5 promoter level mammalian expression atlas. Genome Biol 2015;16:22. doi:10.1186/s13059-014-0560-6.

[35] Forrest ARR, Kawaji H, Rehli M, Kenneth Baillie J, de Hoon MJL, Haberle V, et al. A promoter-level mammalian expression atlas. Nature 2014;507:462–70. doi:10.1038/nature13182.

[36] Javierre BM, Sewitz S, Cairns J, Wingett SW, Vrnai C, Thiecke MJ, et al. Lineage-Specific Genome Architecture Links Enhancers and Non-coding Disease Variants to Target Gene Promoters. Cell 2016;167:1369–1384.e19. doi:10.1016/j.cell.2016.09.037.

[37] Kelly LM, Englmeier U, Lafon I, Sieweke MH, Graf T. MafB is an inducer of monocytic differentiation. EMBO J 2000;19:1987–97. doi:10.1093/emboj/19.9.1987.

[38] Woolf E, Xiao C, Fainaru O, Lotem J, Rosen D, Negreanu V, et al. Runx3 and Runx1 are required for CD8 T cell development during thymopoiesis. Proc Natl Acad Sci U S A 2003;100:7731–6. doi:10.1073/pnas.1232420100.

[39] Ng SWK, Mitchell A, Kennedy JA, Chen WC, McLeod J, Ibrahimova N, et al. A 17- gene stemness score for rapid determination of risk in acute leukaemia. Nature 2016;540:433–7. doi:10.1038/nature20598.

[40] Mori M, Saito K, Ohta Y. ARHGAP22 localizes at endosomes and regulates actin cytoskeleton. PLoS One 2014;9. doi:10.1371/journal.pone.0100271.

[41] Rubin AJ, Barajas BC, Furlan-Magaril M, Lopez-Pajares V, Mumbach MR, Howard I, et al. Lineage-specific dynamic and pre-established enhancer–promoter contacts cooperate in terminal differentiation. Nat Genet 2017. doi:10.1038/ng.3935.

[42] Regev A, Teichmann S, Lander ES, Amit I, Benoist C, Birney E, et al. The Human Cell Atlas. bioRxiv 2017. doi:10.1101/121202.

[43] Shao C, Höfer T. Robust classification of single-cell transcriptome data by nonnegative matrix factorization. Bioinformatics 2017;33:235–42. doi:10.1093/bioinformatics/btw607.

[44] Zhu X, Ching T, Pan X, Weissman S, Garmire L. Detecting heterogeneity in single-cell RNA-Seq data by non-negative matrix factorization. PeerJ Prepr 2016;4:e1839v1. doi:10.7287/peerj.preprints.1839v2.

[45] Darmanis S, Gallant CJ, Marinescu VD, Niklasson M, Segerman A, Flamourakis G, et al. Simultaneous Multiplexed Measurement of RNA and Proteins in Single Cells. Cell Rep 2016;14:380–9. doi:10.1016/j.celrep.2015.12.021.

[46] Genshaft AS, Li S, Gallant CJ, Darmanis S, Prakadan SM, Ziegler CGK, et al. Multiplexed, targeted profiling of single-cell proteomes and transcriptomes in a single reaction. Genome Biol 2016;17:188. doi:10.1186/s13059-016-1045-6.

[47] Clark SJ, Argelaguet R, Kapourani C-A, Stubbs TM, Lee HJ, Krueger F, et al. Joint Profiling Of Chromatin Accessibility, DNA Methylation And Transcription In Single Cells. bioRxiv 2017. doi:10.1101/138685.

[48] Morgan M, Obenchain V, Hester J, Pagès H. SummarizedExperiment: SummarizedExperiment container 2016.

[49] Ouyang Z, Zhou Q, Wong WH. ChIP-Seq of transcription factors predicts absolute and differential gene expression in embryonic stem cells. Proc Natl Acad Sci 2009;106:21521–6. doi:10.1073/pnas.0904863106.

[50] Cairns J, Freire-Pritchett P, Wingett SW, Várnai C, Dimond A, Plagnol V, et al. CHiCAGO: robust detection of DNA looping interactions in Capture Hi-C data. Genome Biol 2016;17:127. doi:10.1186/s13059-016-0992-2.

